# Microplastics clog reproduction in a monkeyflower species

**DOI:** 10.1101/2023.05.21.541634

**Authors:** Gastón O. Carvallo, Valeska Muñoz-Michea

## Abstract

- Plastic debris is a widespread and significant marker of global change, found in all ecosystems and overpassing the total animal biomass on the whole planet. Despite its ubiquity, our knowledge of its potential effects on terrestrial organisms and ecological processes is still limited.
- Recently, a study showed that honey bee *Apis mellifera* transport microplastics (MP; plastic fragments <5 mm) that are added to their bodies. Additionally, a report showed that MP has the potential to reach plant ovules. These findings suggest that pollinators may inadvertently deposit MP on flower stigmas, which could impact the reproductive output of plants.
- We assessed whether small polypropylene fragments (<63 μm) deposited on the stigmas decrease pollen tube development, seed production, seed mass, and germination in the Andean-yellow monkeyflower *Erythranthe lutea* (Phrymaceae).
- Using a mix of hand-pollination treatments and ultrastructure observation, we showed that the MP deposit has a negative impact on seed production and the number of pollen tubes that reached ovaries. However, mass per seed and germination of the resulting seeds were not affected.
- Our findings warn about the deleterious effects of synthetic plastic on a critical ecosystem process, pollination, and suggest that MP could have significant consequences for angiosperms and crop production.

## Introduction

The global mass of synthetic plastic exceeds the biomass of animals on the entire planet (Elhacham *et al*., 2020), resulting in its presence in all environments (Barnes *et al*., 2009; Law & Thompson, 2014; Lavers & Bond, 2017). Microplastics, plastic fragments measuring less than 5 mm (Lim, 2021), are pervasive pollutants that accumulate in various organisms, including humans (Jenner *et al*., 2022). The effects of microplastics (MP hereafter) on terrestrial organisms have been less extensively studied compared to marine organisms (de Souza Machado *et al*., 2018; Baho *et al*., 2021; Huo *et al*., 2022). For instance, in plants MP produces various deleterious effects on metabolism and organ development (Qi *et al*., 2018; Dong *et al*., 2021; Sun *et al*., 2021; Zantis *et al*., 2023), but fewer studies have assessed their impacts on terrestrial ecosystem processes involving plants (but see Abdolahpur Monikh *et al*., 2022; Zhu *et al*., 2022).

Biotic pollination, the animal-mediated transfer of pollen grains among plants, is a crucial ecosystem process because approximately 87.5% of angiosperms (Ollerton *et al*., 2011) and around 85% of crops (Klein *et al*., 2007) rely on animals for reproduction and seed generation. Biotic pollination might be severely impacted by MP due to the fact that pollinators, typically arthropods, contact with soil and water basins where MP accumulate (Balzani *et al*., 2022). Indeed, the honey bee *Apis mellifera*, which is one of the most conspicuous and important pollinators (Hung *et al*., 2018), has been found to transport MP (Edo *et al*., 2021) that potentially alter their foraging behavior (Deng *et al*., 2021) and their effectiveness as pollinator vectors (Balzani *et al*., 2022). Furthermore, MP have been detected in honey bee-derived products, including honey (Diaz-Basantes *et al*., 2020; Al Naggar *et al*., 2023).

In plants, studies on the effects of MP have primarily focused on their impact on vegetative facets (Rillig *et al*., 2019). For instance, they have been found to decrease photosynthesis efficiency (Dong *et al*., 2021) and affect plant growth and biomass (Lian *et al*., 2021; Zantis *et al*., 2023). Microplastics are mainly taken up by plants from soils and leaves (via aerosols) and are then translocated to different tissues (Qi *et al*., 2018; Sun *et al*., 2020; Urbina *et al*., 2020). Regarding reproductive effects, it has been suggested that MP-polluted pollinators could inadvertently deposit plastic fragments on flower stigmas during foraging visits (de Souza Machado *et al*., 2018; Oliveira *et al*., 2019). However, to the best of our knowledge, no studies have reported on this specific issue. The only existing study that explored the relationship between MP and plant reproduction conducted an experiment in which latex polystyrene beads measuring 6 μm were artificially deposited on stigmas to assess the role of the stylar matrix in pollen tube development (Sanders & Lord, 1989). This pioneering study demonstrated that latex beads were translocated from the stigma to ovaries, reaching the ovules. However, the study did not quantify the impacts on fertilization and seed development (Sanders & Lord, 1989).

Despite the increasing presence of MP in terrestrial ecosystems (Barnes *et al*., 2009), there is currently a lack of robust evidence to determine their impacts on the pollination process and plant reproduction. Recognizing this critical knowledge gap, we conducted a study to assess the effects of polypropylene fragments on seed production. Specifically, we deposited fragments of MP (<63 μm) on the stigmas of the Andean-yellow monkeyflower *Erythranthe lutea* (Phrymaceae), which is widely used as a model species in ecological and evolutionary studies (Yuan, 2019). Our main objective was to examine whether the presence of MP on the stigmas inhibits pollen tube development, analogous to the effect of heterospecific pollen (Wilcock & Neiland, 2002; Broz & Bedinger, 2021), thereby clogging the stigmatic surface and reducing plant reproductive output. We discuss our findings and their implications for pollination of both wild and cultivated plants, taking in consideration the limitations of our study. Finally, we propose future research directions to address knowledge gaps concerning the physiological and ecological consequences of plastic debris on plant-pollinator interactions and plant reproduction, aiming to understand this emerging environmental health threat.

## Materials and methods

### Studied plant specimens

The Andean-yellow monkeyflower, *Erythranthe lutea* (L.) G.L. Nesom (Phrymaceae), is a native species from Chile, which produces abundant large flowers per individual (von Bohlen, 1995). This species is self-compatible and exhibits some degree of autonomous reproduction (Carvallo & Medel, 2010). For our study, we utilized a first-generation of selfing individuals derived from seeds collected in 2021 from Farellones (individual lut039), Laguna del Maule (lut061) and Ralco (lut019) (Supplementary Material, Table S1). These seeds were germinated in December 2021 after being refrigerated at 4° C for 10 days in 245 cm^3^ pots filled with a commercial substrate (Anasac, Santiago). Then, the pots were transferred to growth chambers (SRI21D-2; Sheldon Manufacturing Inc., Cornelius, Oregon) and maintained under a light-dark cycle of 18/6 hours at 21° and 16° C, respectively. Once the plants reached approximately 30 mm in height, they were individually potted and kept in the growth chamber. Plants were watered every two to three days with 50 mL of water and fertilized using BioBloom (BioGrow, Bizkaia, Spain) according to the manufacturer’s recommendations (2 mL per liter of water every two weeks). Throughout the experiments, plants were kept under these controlled conditions until they reached the flowering stage, and all the subsequent experiments were conducted inside the growth chambers to minimize any potential interference from external sources of pollen or plastic.

### Microplastic provisioning

We obtained transparent polypropylene flakes from a semi-industrial plastic recycling plant (Revuelta, Valparaíso). We chose this kind of plastic to provide a more realistic scenario for our assays, in line with recommendations by some authors ((Zimmermann *et al*., 2021; Zantis *et al*., 2023). Polypropylene is commonly abandoned, often in illegal landfills, after being extensively used as greenhouse covers and hoses in the agro-industry. Clean and dry polypropylene flakes, approximately 10 mm in size, were obtained using a shredder. The flakes were transported to the lab and soaked in a solution of commercial dishwashing liquid and a 1% chlorine bleach solution for 24 h. Then, the flakes were rinsed 4-5 times with tap water, with the final rinse conducted using distilled water. A volume of 300 ml of dry plastic flakes was mixed with an equal amount of distilled water, and this solution was blended in a juice blender machine (Oster, Milwaukee, WI) for 10 min. The mixture was then dried at 50° C in a dry oven for six consecutive days. The resulting material was sifted using a set of sieves (Humboldt, Elgin, IL) with gradations of 1000, 500, 250, 125 and 63 μm. The remaining dust that passed through the 63 μm sieve, approximately 10 g, was used in our assays and considered as MP. To visualize their size and shape and compare them with pollen from the studied plant, we performed observations of the obtained plastic dust under a light microscope (Supplementary Material, Fig. S1a).

### Pollination treatments and reproductive output

As the plants reached the bloom stage, we tagged flower buds. On the second day of anthesis, we applied one of the following hand-pollination treatments to each flower, following a previously established random sequence. (i) Hand self-pollination: using fine forceps, we carefully picked out one anther, deposited it in a mini glass tube, and stirred it with a wooden toothpick. The pollen that covered the tip of a toothpick was then deposited on the stigma surface of the focal flower. (ii) Plastic-pollen mix: an anther was extracted from the focal flower and mixed with an equivalent amount of microplastic dust within a mini glass tube. The mixture that covered the tip of the toothpick was then deposited on stigma. (iii) Plastic-then-pollen: in this treatment, we deposited an amount of plastic dust that fit the top of a toothpick onto the stigmas. Then, when stigmas opened (*E. lutea* has touch-sensitive stigmas; Carvallo & Medel, 2010), we deposited pollen from the same focal flower onto the stigma using the tip of a toothpick. (iv) Pure plastic: an amount of plastic dust equivalent to the tip of a toothpick was directly deposited onto stigmas. In all of the aforementioned treatments, flowers were emasculated by removing the anthers. Additionally, we studied a group of unmanipulated flowers as control. A total of 153 flowers from 10 individuals were included in the study. Between fifteen and twenty days after the treatments, we recovered the capsules, which were then individually placed in paper bags and dried 48 h at 40° C in a dry oven. The seeds from each capsule were counted under a stereomicroscope, and the total mass of seeds per fruit was measured using an ultra-analytic balance (M500P, Sartorius, Goettingen). Based on the number of seeds per fruit and their weight, we estimated the average mass of each seed.

### Seed germination

We assessed whether seeds obtained from pollination treatments described in the previous section differed in their germination levels. From each fruit, we randomly selected ten seeds, which were then placed in 50 mm diameter glass Petri dishes using commercial soil as substrate (Anasac, Santiago). At the start of the assay, the seeds were watered with 10 mL of distilled water. We included 15 replicates for each treatment. However, due to the lower number of fruits obtained in some pollination treatments, we reduced the number of replicates for those treatments (plastic-pollen mix: 10 replicates; plastic-then-pollen: 5 replicates). Since the pure-plastic treatment resulted in almost no seed production, it was not included in this part of the study. The Petri dishes with the sown seeds were placed in a fridge at 4° C for ten days and then transferred to a growth chamber, maintaining the previously described conditions (18/6 hours of light and dark at 21°/16° C, respectively). Every two-three days, we monitored the number of germinated seeds in each dish. Seeds were considered germinated when cotyledons emerged. The germination process was monitored for a duration of 30 days.

### Ultrastructure observation and pollen tubes

We employed two approaches to assess the impact of MP on stigmas and their potential effects on pollen tube development: fluorescent light microscopy (FLM) and scanning electron microscopy (SEM). For FLM, we prepared five flowers for each treatment described above (self-hand pollination, plastic-pollen mix, plastic-then-pollen, and unmanipulated), with the exception of the pure plastic treatment (N = 4 flowers). Forty-eight hours after applying treatments, we extracted and fixed the complete gynaeceum in Farmer’s solution for 24 hours (Thompson, 2016). Then, we rinsed the tissue in distilled water for 10 minutes and soaked it in an 8 M NaOH solution for 24 hours to soften the tissues. After softening, the tissue was washed again in distilled water and stained overnight in a 0.1% aniline blue solution (Oneal *et al*., 2016). We observed samples using an Olympus CX31 microscope (Shinjuku, Tokio) equipped with a fluorescence lamp. In these samples, we counted the number of pollen tubes reaching the upper portion of the ovary.

For SEM, we examined stigmas from the five treatments (self hand-pollination, plastic-pollen mix, plastic-then-pollen, pure plastic and control). We mounted stigmas samples on stubs using double-sided carbon adhesive tape. Non-metalized samples were then observed under partial vacuum using a HITACHI SU 3500 SEM (Chiyoda, Tokio, Japan) at an accelerating voltage of 20 kV. The obtained images provided qualitative information regarding the distribution of MP, pollen and pollen tubes on stigmas.

### Statistical analysis

To determine the effects of treatments on the number of seeds and seed mass, we fitted a generalized linear mixed model (GLMM) using restricted maximum likelihood REML with the ‘*lmer’* function from the ‘lme4’ package in R 4.1.1 (R Core Team, 2017). Treatments were included as a fixed factor, and individual plants (N = 10 plants studied) were considered as a random factor. Fixed effects were assessed using type III analysis of variance with Satterthwaite’s method implemented with the ‘*lmerTest’* function (Kuznetsova *et al*., 2017). To analyze the effects of pollination treatments on seed germination, we employed two approaches. Firstly, we compared the cumulative distribution function curves among treatments using nonparametric time-to-event fitting to estimate the nonparametric maximum likelihood estimator (NPMLE) following the procedures outlined in (Onofri *et al*., 2022). This analysis was performed using the ‘*drmte*()’ and ‘*compCDF*()’ functions from the ‘*drcte’* package in R (Onofri *et al*., 2022). Secondly, we used a more traditional approach by comparing the cumulative proportion of germinated seeds at the end of the germination assay, employing a generalized linear model (GLM) with treatments as a fixed factor. Furthermore, we compared the number of pollen tubes among pollination treatments based on the FLM images using an analysis of variance (ANOVA) with Satterthwaite’s method, also implemented with the ‘*lmerTest’* function (Kuznetsova *et al*., 2017). The dataset and script used in this study are available on GitHub (https://github.com/gcarvallob/ErythrantheMicroplastics2023).

## Results

The presence of MP in pollination treatments resulted in a decrease in seed production (Fig. 1a; Supplementary Material, Table S2a), while the seed mass was not significantly affected (Fig. 1b; Supplementary Material Table, S2b). Specifically, the pollen-plastic mix treatment yielded 7.3 ± 2.5 seeds (mean ± s.e., N = 30 fruits analyzed), and seed production was even lower in the plastic-then-pollen treatment (3.3 ± 1.1 seeds, N = 29 fruits). Flowers that were exposed to plastic alone (without pollen) did not produce seeds, except for two fruits (0.7 ± 0.6 seeds; N = 32 fruits). These results contrasted with the hand self-pollinated treatment (198.5 ± 43.8 seeds; N = 30 fruits) and the control group (36.9 ± 10.4 seeds; N = 33 fruits). ANOVA using Satterthwaite’s method showed that seed production significantly differed among the treatments (d.f. = 4; *F* = 18.028, *P*<0.001). The model’s total explanatory power was substantial (R^2^_conditional_ = 0.33), and the effect of individual plants on seed production was marginal (Supplementary Material, Fig. S2a). The fixed effects alone accounted for a greater proportion of the explanatory power of the model (R^2^_marginal_ = 0.32).

**Figure 1.**
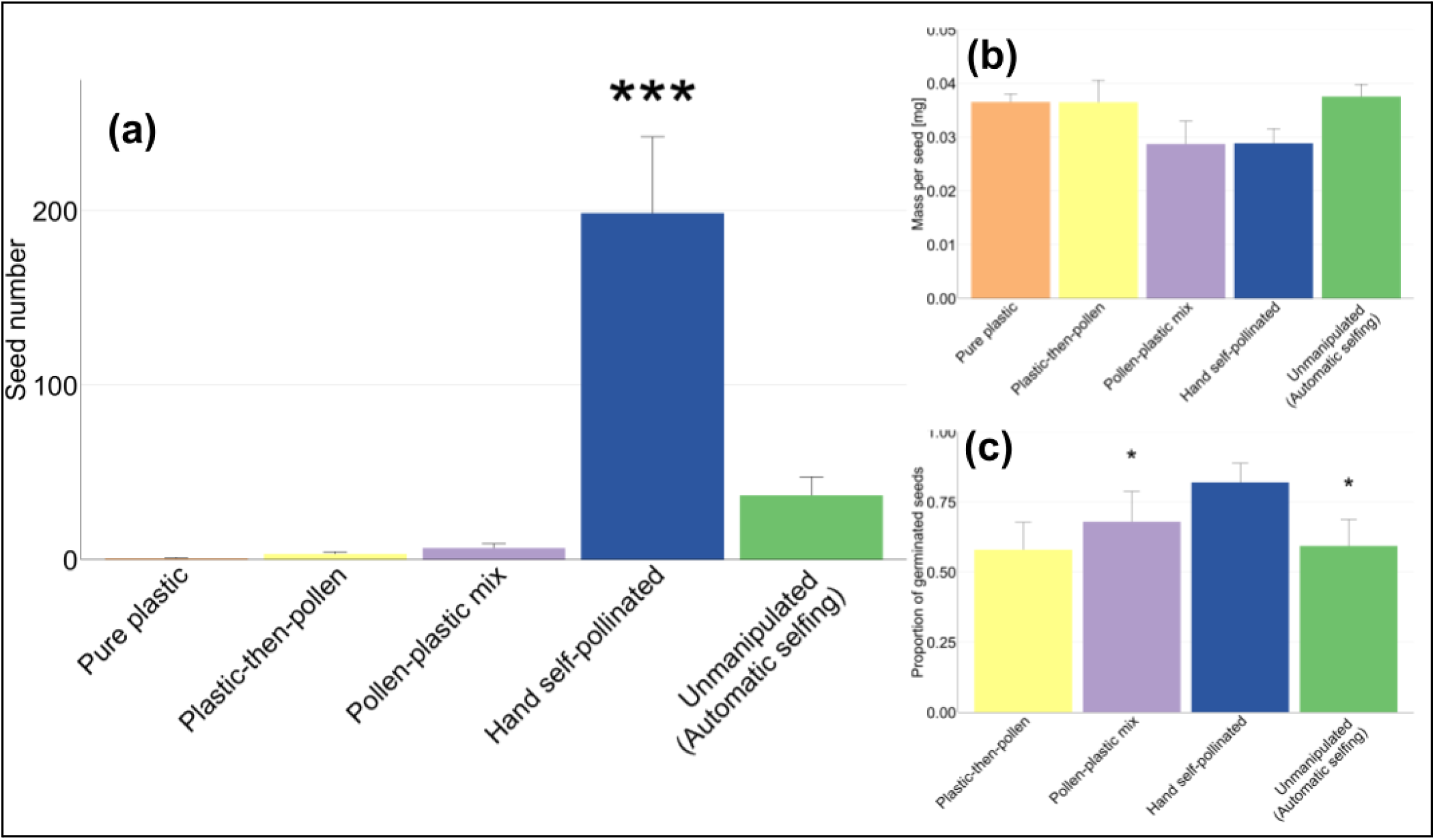
Effects of microplastic on (a) seed production, (b) seed mass, and (c) seed germination of *E. lutea* (mean ± se) We assessed the effects of polypropylene fragments (<63 μm) mixed with an equivalent amount of pollen (pollen-plastic mix treatment) and the sequential deposit of plastic followed by pollen (plastic-then-pollen treatment). Additionally, we included flowers “pollinated” with plastic alone (pure plastic treatment) and a group of unmanipulated (control) flowers. Variables that significantly differed from the intercept are depicted with asterisks (refer to Supplementary Material, Table S2a). ***P<0.001 after GLMM.

We estimated a seed mass of 0.03 ± 0.01 mg per seed (range: 0.00 -0.07 mg; based on 84 analyzed fruits). Seed mass did not vary significantly among the different pollination treatments (Fig. 1b), as indicated by ANOVA (d.f. = 4; *F* = 0.583, *P* = 0.675; parameters estimated using Satterthwaite’s method). The effect of individual plants on the model was substantial (Supplementary Material, Fig. S2a), while the contribution of fixed effects alone was minimal (R^2^_marginal_ = 0.02) to the overall explanatory power of the model (R^2^_conditional_ = 0.45). Regarding seed germination, we did not observe significant differences among treatments in terms of germination curves (T_observed_ = 0.477; *P* = 0.645; Supplementary Material, Fig. S3) and germination at the end of the assay (d.f. = 3; *F* = 2.463; *P* = 0.077). Overall, we observed a germination proportion of 0.69 ± 0.05 seeds (N = 45 plates), with a higher proportion in the seeds obtained from the hand self-pollinated treatment (0.82 ± 0.07 seeds germinated; N = 15 plates; Fig. 1c) and a lower proportion in the plastic-then-pollen treatment (0.58 ± 0.09 seeds germinated; N = 5 plates).

Hand self-pollinated flowers developed a higher number of pollen tubes (235 ± 77 pollen tubes, N = 5 flowers) compared to treatments that received plastic (Fig. 2). The results of ANOVA with Satterthwaite’s method indicated a significant difference among the treatments (d.f. = 2; *F* = 5.813, *P* = 0.018). Notably, the pure plastic and control (unmanipulated) treatments did not show any pollen tube development (Fig. 2b). Scanning electron microscopy (SEM) analysis of stigmas revealed potential blockage of stigma papillae by MP, which could impact the germination of pollen tubes (Fig. 3). The presence of plastic fragments appeared to act as a barrier to pollen grain germination. In the plastic-then-pollen treatment, fewer pollen tubes were observed, and the majority of the pollen grains were deposited on the plastic layer (Fig. 3d, Fig. 3i). In the pollen-plastic mix treatment, we observed a higher number of MP fragments and pollen tubes development (Fig. 3e, Fig. 3g). Both the pollen grains and pollen tubes in flowers that received MP did not show any irregular morphology compared to the control treatments (Fig. 3). Overall, the treatments that received plastic displayed size variation in MP fragments, including tiny fragments surrounding pollen and pollen tubes, suggesting the presence of nanoplastics (Fig. 3g Fig. 3i-j).

**Figure 2.**
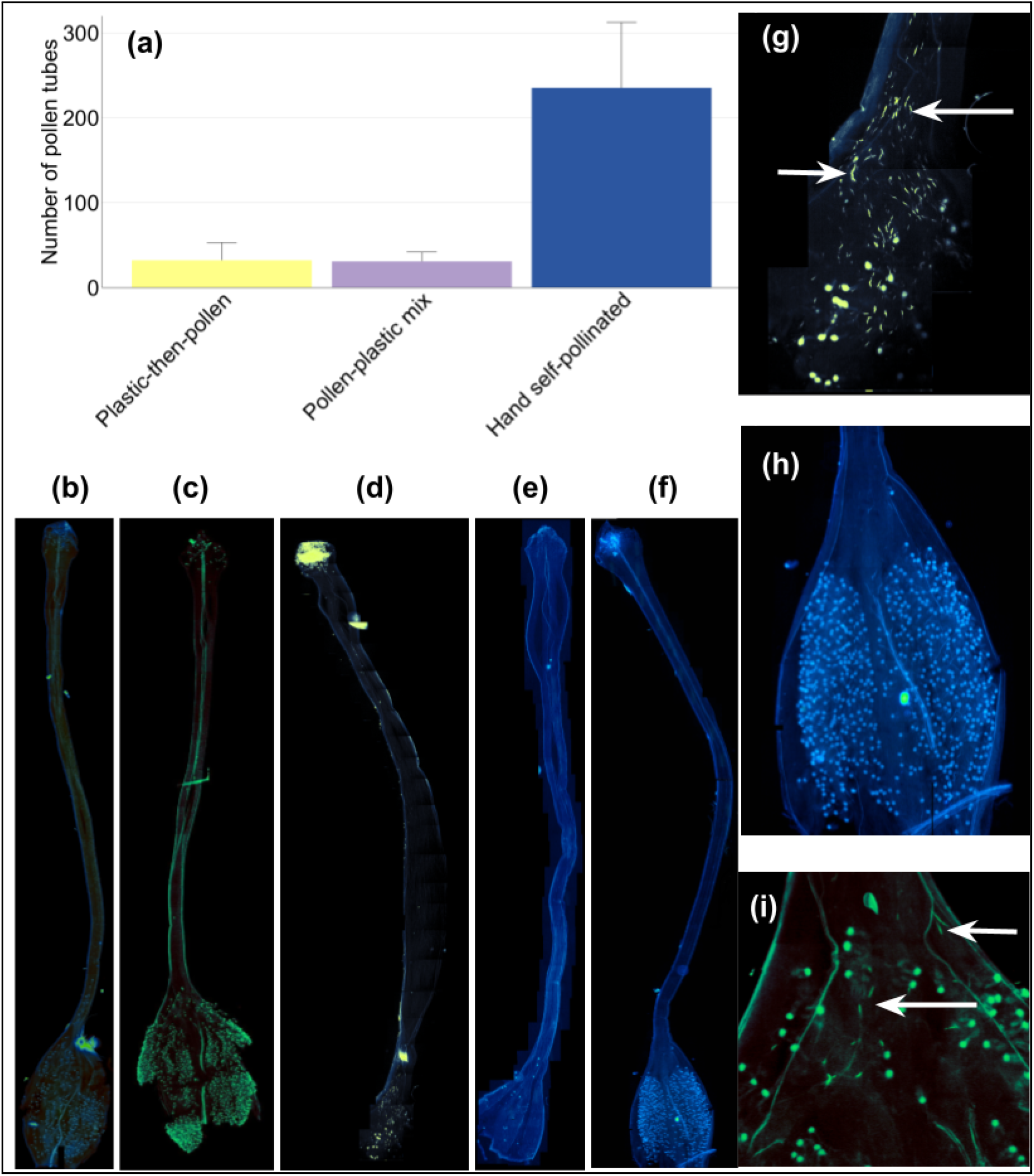
Effects of microplastics on pollen tube development *in E. lutea*. **(a)** Number of pollen tubes (mean ± se) in treatments that received plastic and pollen. Treatments that received only plastic and unmanipulated flowers (control) were not included in this plot as they did not produce pollen tubes. **(b-f)** Microphotographs of *E. lutea* gynoecium, and **(g-i)** detailed images of ovaries. **(b)** Plastic-then-pollen, **(c-i)** pollen-plastic mix, **(d-g)** hand self-pollinated, **(e)** control, **(f-h)** pure plastic. White arrows indicate some of the observed pollen tubes. Images were obtained by merging a sequence of pictures taken under a microscope with a fluorescence lamp (10x).

**Fig. 3.**
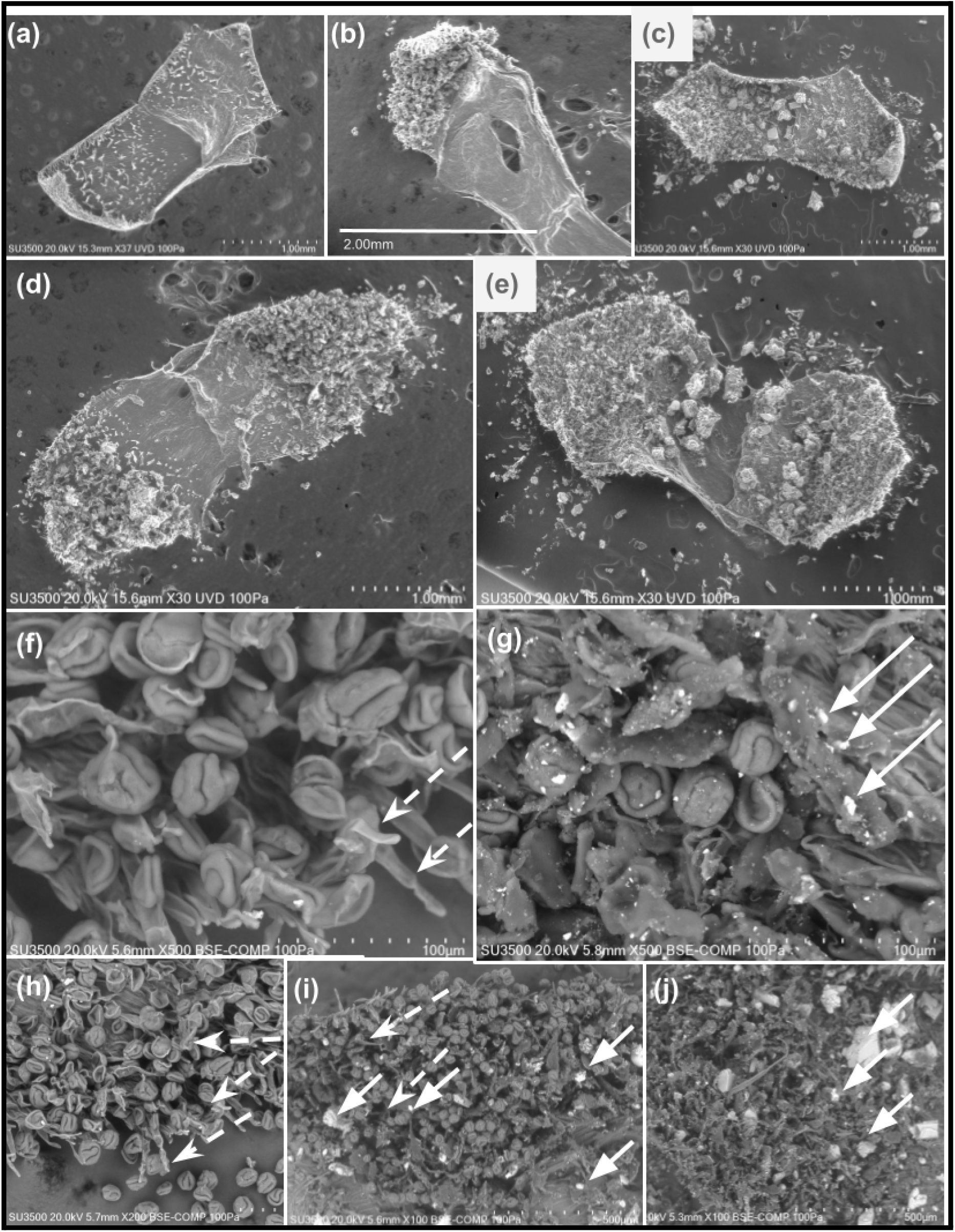
SEM microphotographs of stigmas under different plastic deposit treatments. Stigmas were collected from two-day old flowers. (a) Unmanipulated flower showing prominent stigma papillae. (b, f, h) Hand self-pollinated flowers. (c, j) “Hand-pollinated” using plastic. (d, i) Plastic-then-pollen. (e,g) Pollen-plastic mix. *Continuous white arrows* depict plastic fragments in high magnification images. *Dotted white arrows* depict pollen tubes.

## Discussion

In this study, our objective was to investigate the potential impact of MP deposition on stigmas on the reproductive output of plants. Our findings revealed a significant decrease in seed number after following MP deposition in the Andean-yellow monkeyflower. This astonishing result highlights the potential threat posed by MP to terrestrial organisms involved in complex interactions, such as pollination. The implication of MP-induced reduction in reproductive success extends beyond our study system and could have implications for crop production, particularly in plant species inhabiting plastic-polluted environments, which is a common occurrence in certain regions (Corradini *et al*., 2019). Previous research has shown that the presence of MP in substrates can lead to significant changes in vegetative traits, including reduced aerial and root growth, biomass accumulation, germination, and alterations in metabolism (Zantis *et al*., 2023). While there have been advances in understanding the effects of MP on vegetative characteristics, our study provides the first evidence demonstrating the impacts of MP on plant reproduction.

Although MP deposited on stigmas can potentially affect various fitness components, we did not detect any impacts on seed mass and germination in our study. However, we did find a reduced number of pollen tubes in flowers treated with plastic, indicating that MP may mechanically obstruct the passage of pollen tubes, similar to the effect of heterospecific pollen (Wilcock & Neiland, 2002). Nevertheless, it is important to consider other characteristics of MP, such as its chemical structure, electric charge, and the presence of additives, which could potentially influence pollen tube development through chemical inhibition. For example, residues from other types of pollutants, such as graphene, have been shown to impact pollen tube development in *Cucurbita pepo* (Candotto Carniel et al., 2018, 2020). Therefore, MP may have both mechanical and chemical impacts on pollen adhesion to stigmas and pollen tube development. Additionally, it is necessary to incorporate methods that assess the effects of stigma accumulated layers of plastic on pollen tube development. This is particularly crucial considering that plants are exposed to multiple successive events of MP deposition.

Some results obtained in our study may seem counterintuitive, but they can be largely attributed to the characteristics of the model species used. For example, the production of seeds in the “only plastic” treatment (two fruits) may result from delayed selfing, which is an assurance reproductive mechanism found in some *Erythranthe* species (Dole, 1992; Vickery, 2008). On the other hand, the presence of pollen tubes and seed formation in the plastic-them-pollen treatment may be explained by some degree of selfing that occurred before the manipulative treatments were applied. Since we performed hand-pollination in two-day-old flowers, it is possible that there was sufficient time for automatic selfing to take place. Alternatively, in this treatment pollen grains might be capable of partially crossing the plastic layer deposited on the stigmas, as supported by our observations of pollen tubes under FLM and SEM. Another counterintuitive result is the absence of pollen tubes in the control (unmanipulated) group, despite the production of seeds. We believe that this result can be attributed to the fact that the control group had ample time to develop selfing assurance (Dole, 1992; Vickery, 2008) before the pollination experiment was performed, which facilitated seed production. However, the absence of detectable pollen tubes in the control group may be due to the fact that flowers were collected only 48 hours after anthesis, which might have been an insufficient amount of time to observe pollen tubes through selfing assurance. The use of a standard and well-established methodology in this study ensures a high level of replicability for our experiments. This not only enhances the reliability of the findings but also provides an opportunity to explore additional variables that were not included in this study, thereby paving the way for future experimental designs.

Our study raises several research questions that are crucial to address in order to determine the effect of plastic debris on pollination and plant reproduction. Several authors have suggested that MP are likely to interfere with pollination (de Souza Machado et al., 2018; Oliveira et al., 2019; Deng et al., 2021). However, in order to identify the impacts of plastic debris on pollination, it is important to consider other variables such as fragment size and shape, MP concentration, as well as chemical composition and presence of plastic additives. In the following sections, we provide an overview of some of these factors, recognizing both the direct and indirect impacts of MP on animals and plants involved in pollination (Fig. 4).

**Fig. 4.**
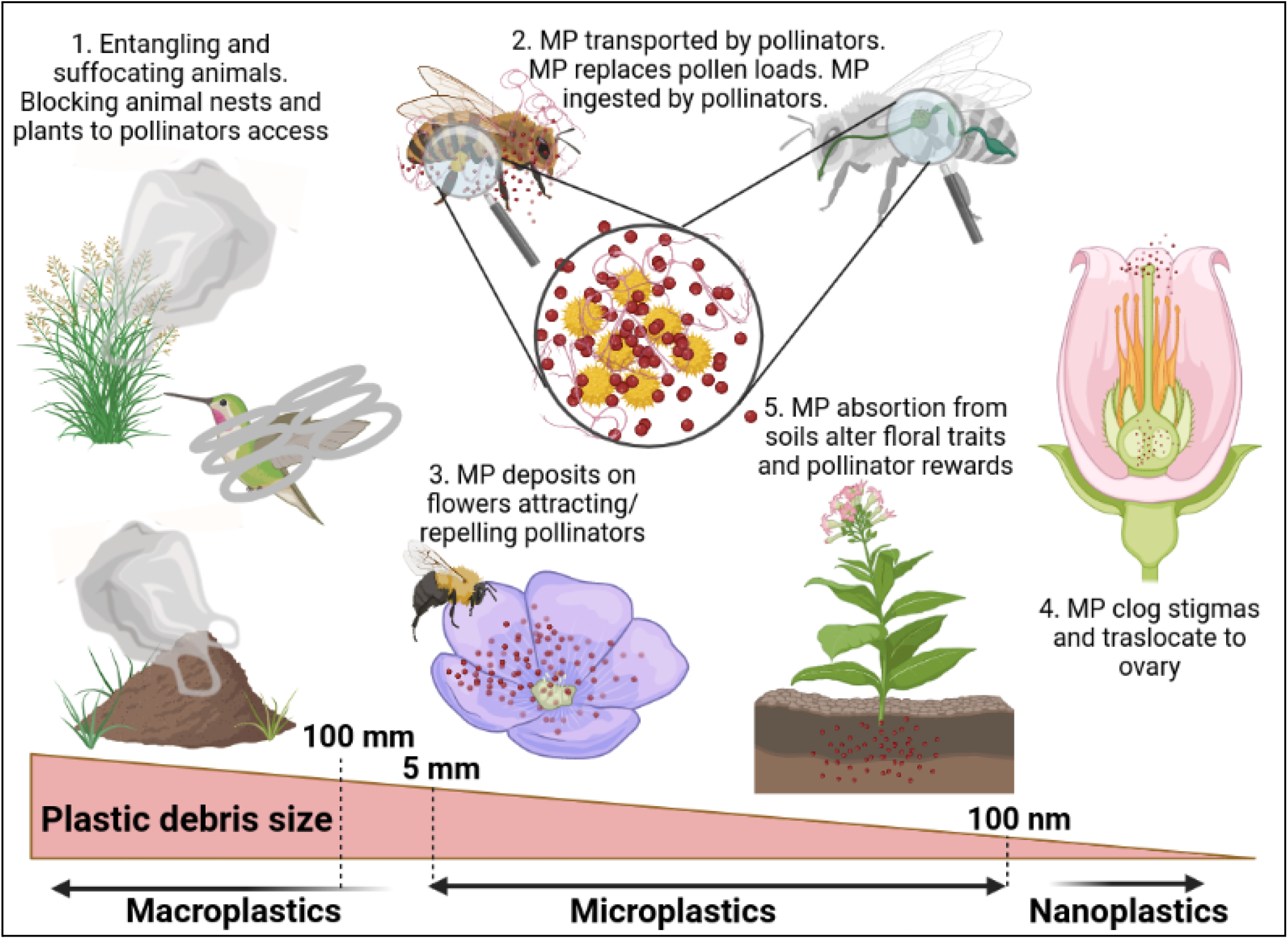
Potential direct and indirect effects of plastic debris on pollination. The numbers correspond to the compilation details provided in the text. MP: microplastics. Figure created with BioRender.com

### 1. Macroplastics and pollination

Macroplastics are large plastic debris that are visible to the naked eye, typically measuring more than 100 millimeters (referred to as mega-debris by some authors, see (Barnes *et al*., 2009). These include items such as plastic bags, bottles, and packaging materials that are commonly found in the ocean and on land. Macroplastics can pose harm to wildlife by entangling and suffocating animals (Barnes *et al*., 2009) or by being ingested, which can cause blockages or physical damage to the digestive system of pollinators (Nelms *et al*., 2015; Lusher & Hernandez-Milian, 2018). Although no studies have directly connected macroplastics and pollination, plastic debris can have indirect effects by affecting larger pollinators (i.e., suffocating vertebrates). Macroplastics can also obstruct nest entrances for pollinators or impede pollinator access to flowers (Fig. 4). The obstruction caused by plastics could be a significant threat to plant species with restricted distributions, particularly those inhabiting areas near polluted sites (i.e., illegal landfills). For instance, in the South American mediterranean-type ecoregion, plants, especially cacti species, surrounding illegal landfills have been observed with flowers covered in macroplastics (Carvallo, pers. obs.). This situation could be exacerbated by certain plant traits that facilitate the adhesion of plastics on plants, such as spines or sticky leaves.

### 2. Pollinator-mediated microplastics transport

Arthropods inadvertently pick up small plastic particles and can transport them from one location to another (Edo *et al*., 2021; Balzani *et al*., 2022), allowing for their potential spread over long distances. For instance, *Bombus terrestris* has the capacity to cover distances of up 2.5 km, spanning an area of 0.25-4 ha (Hagen *et al*., 2011). Pollinators can acquire MP through various means, primarily from soils and water sources (Balzani *et al*., 2022). The negative electric charge of MP, similar to that of pollen grains, may attract them to pollinators, which are positively charged (Clarke *et al*., 2013). Additionally, MP could potentially substitute or be incorporated into the pollen loads carried by pollinators. Understanding the extent and mechanisms of animal-mediated MP transport is crucial for assessing their spread in the environment. Investigating the movement of different types and sizes of MP fragments and their potential impacts on foraging range of pollinators remains an open question.

### 3. Microplastics and pollinators’ health

One of the rapidly growing areas of research focuses on assessing the effects of MP on the health and the behavior of honey bees (Wang *et al*., 2021; Deng *et al*., 2021; Balzani *et al*., 2022; Al Naggar *et al*., 2023). Pollinators can ingest MP through various routes, including contaminated pollen, nectar, or direct contact with contaminated surfaces (Deng *et al*., 2021). In honey bees, the ingestion of MP increases their vulnerability to viral infections, with small fragments even translocating to the haemolymph (Deng *et al*., 2021). The accumulation of MP in tissues could potentially impact pollinator behavior, as observed in *A. mellifera* (Balzani *et al*., 2022). This could result in suboptimal foraging behavior, limiting their movement range or the time spent on flowers. Furthermore, MP has been suggested to act as pathogen vectors among honey bees (Al Naggar *et al*., 2023), although their own microbiome helps protect them from MP ingestion (Wang *et al*., 2021). However, the full extent of MP’ impact on pollinators’ health remains unknown. For instance, MP may damage the digestive system, leading to reduced feeding efficiency and nutrient absorption (Oliveira *et al*., 2019; Deng *et al*., 2021). Also, plastic additives may be toxic to arthropods, affecting various stages of their development (Hahladakis *et al*., 2018). It is crucial to investigate how animals traits, such as body size, thermoregulation mechanisms (in ecto- and endotherms pollinators), pilosity density, and characteristics of corbiculae and scopae characteristics (which may act as MP baskets), can modulate the health response of pollinators to microplastics.

### 4. Microplastic deposition in flowers

Fragments deposited on flowers can alter the cues for pollinator visits (Fig. 4), especially visual cues, as plastic is an attractive item for animals (Santos *et al*., 2021). Pollinators may avoid plastic fragments if they perceive them as potential threats, such as predators (Vieira *et al*., 2017), or if they alter corolla brightness and iridescence, which are crucial cues for pollinator attraction (Whitney *et al*., 2009; Hempel de Ibarra & Holtze, 2022). Plastic fragments could potentially attract pollinators to flowers, especially if the MP fragments emit similar wavelengths of light as flowers. For instance, in the water column, some fish have shown a preference for certain colors of MP (Ory *et al*., 2017). Therefore, plastic fragments can have important implications as attractive or adverse cues for pollinators (Fig. 4), as they can affect their behavior and efficiency, potentially reducing the time that animals spend on flowers.

A second implication of MP deposition on flowers is the impact on plant reproduction, as reported in this study. However, our current knowledge is limited to only one species (*E. lutea*), one type of plastic (polypropylene), and one range of fragment sizes (<63 μm). To the best of our knowledge, only one study has reported the translocation of nano-particles (latex beads of 6 μm) to ovaries, reaching ovules, although they did not assess the effects on seed production (Sanders & Lord, 1989). Our observations using SEM, which revealed the presence of small plastic fragments on the stigmas of polluted plants, motivate further investigation into whether nanoplastics can translocate to the ovary and reach ovules, in order to determine if the phenomenon described by (Sanders & Lord, 1989) applies to other plant species.

### 5. Alteration of flower traits mediated by the absorption of microplastics from soils

Soils acts as sinks for MP (Rillig *et al*., 2021; Zantis *et al*., 2023), which can modify the growth and physiology of plants (Zantis *et al*., 2023). For example, the presence of MP in the substrate can reduce the aerial and root biomass of plants (Sun *et al*., 2020) and alter the concentration of pigments, particularly carotenoids (Zantis *et al*., 2023). This primarily occurs because MP impedes water and nutrient uptake (Urbina *et al*., 2020; Zantis *et al*., 2023). Additionally, MP can translocate and accumulate in the aerial tissues of plants (Sun *et al*., 2020; Spanò *et al*., 2022; Zhu *et al*., 2022), thereby altering the length of vegetative structures such as stems and leaves (Zantis *et al*., 2023).

Despite the relatively well-documented impacts of MP on plant growth and vegetative traits after following soil absorption, there is a significant gap in our understanding of how these contaminants might affect flower traits that serve as cues or rewards for pollinators. Traits such as phenology, flower number, corolla size, flower pigment concentrations, nectar volume and their concentration, as well as pollen production, could be modified when plants absorb and translocate MP. This situation could indirectly alter the attractiveness of plants to pollinators, especially in hydrophilic species, such as those used in our study, which could uptake MP directly from water. However, this phenomenon remains largely unexplored and warrants further investigation.

### 6. Increasing the realism of MP in pollination studies

Plastic production and the deposition of MP is an ongoing process that will continue in the coming years (Lebreton & Andrady, 2019). This scenario requires studies that anticipate the potential impacts that living beings will experience. While MP has not yet been identified as a threat to pollinators and pollination (Brown *et al*., 2016), it is important to enhance the realism and reliability of studies on MP and their impact on pollination. Incorporating a range of field studies examining a greater number of species, both pollinators and plants, is crucial to understand how MP moves in the environment and how they alter both natural and agricultural ecosystems. As MP increases in the airborne environment (Trainic *et al*., 2020; Shao *et al*., 2022), small fragments could also affect anemophilous pollination. Researchers should collect more comprehensive environmental information about MP (Trainic *et al*., 2020), including their concentrations in soils, air, and plants. By doing so, it will be possible to establish the relationships between MP concentrations in different environmental contexts (i.e. geographic, climatic, and disturbance gradients) and their concentrations in pollinators and plants. Incorporating these factors into studies of MP and pollination will be essential for advancing our knowledge and developing effective strategies to mitigate their harmful effects.

## Conclusion

Our results demonstrate a significant decrease in seed number following the deposition of MP on stigmas, indicating that this material poses a potential threat to the reproductive success of plants. Although MP does not appear to alter seed mass and germination, our findings suggest that MP can mechanically affect pollen adhesion to stigmas and impede the development of pollen tubes. These conditions are particularly relevant in plastic-polluted areas. This study raises several research questions that need to be addressed to determine the effects of plastic debris on pollination and plant reproduction at different levels. Overall, our study provides the initial evidence highlighting the impacts of MP on plant reproduction, serving as a warning in our current plastic-polluted world.

## Acknowledgements

This study was partially funded by Centro Basal IEB ANID [grant FB210006].

## CRediT authorship contribution statement

**G.O.C**.: conceptualization, data curation, formal analysis, funding acquisition, investigation, methodology, resources, supervision, validation, visualization, writing -original draft, writing -review & editing. **V.M-M**.: data curation, investigation, validation.

## Supplementary Material

**Table S1.**
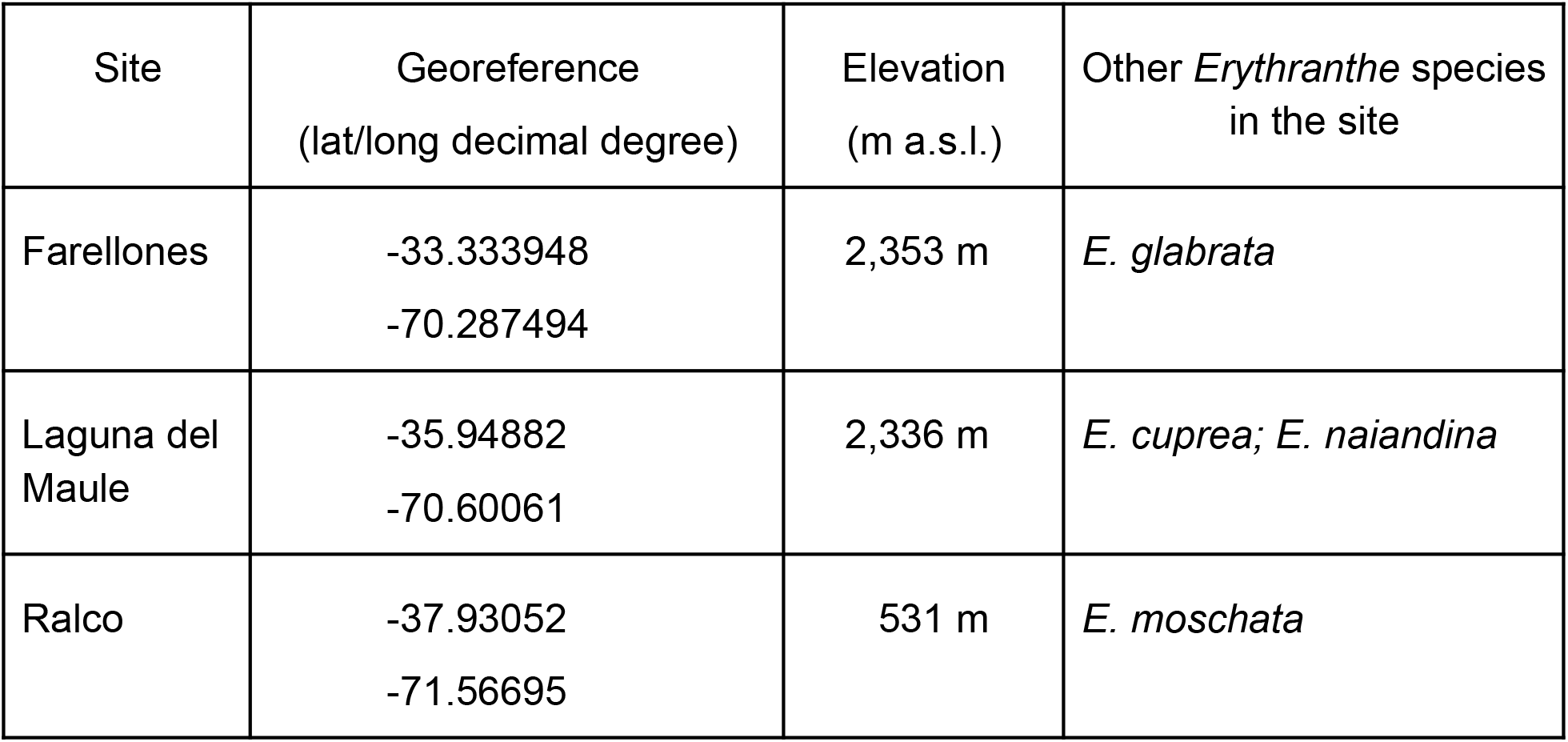
Sites from which studied plants were obtained

**Table S2.**
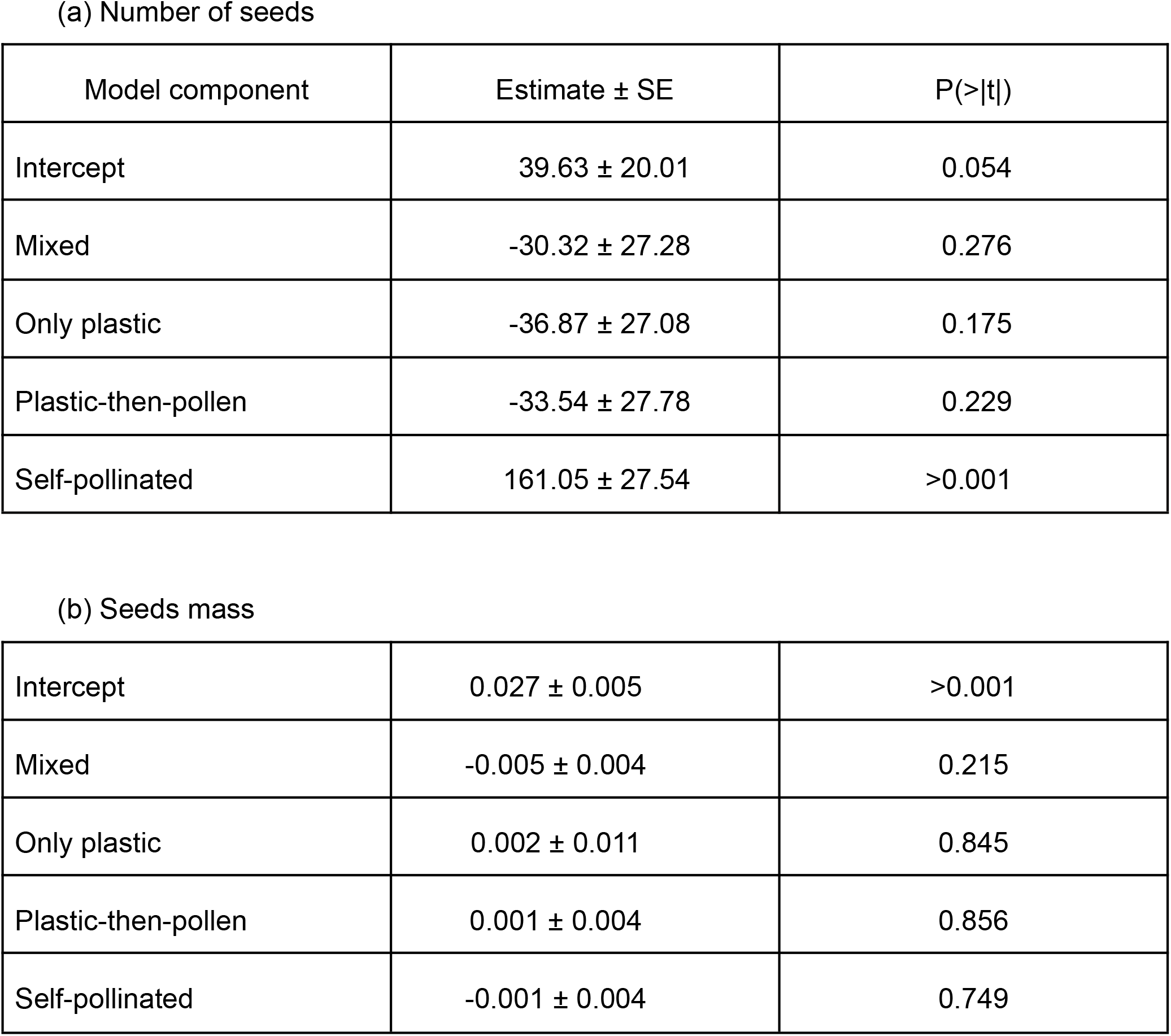
Effect of pollination treatments (fixed factor) on **(a)** the number of seeds per fruit,and **(b)** seed mass. Parameter estimators were obtained with a GLMM. SE: standard error.

**Figure S1.**
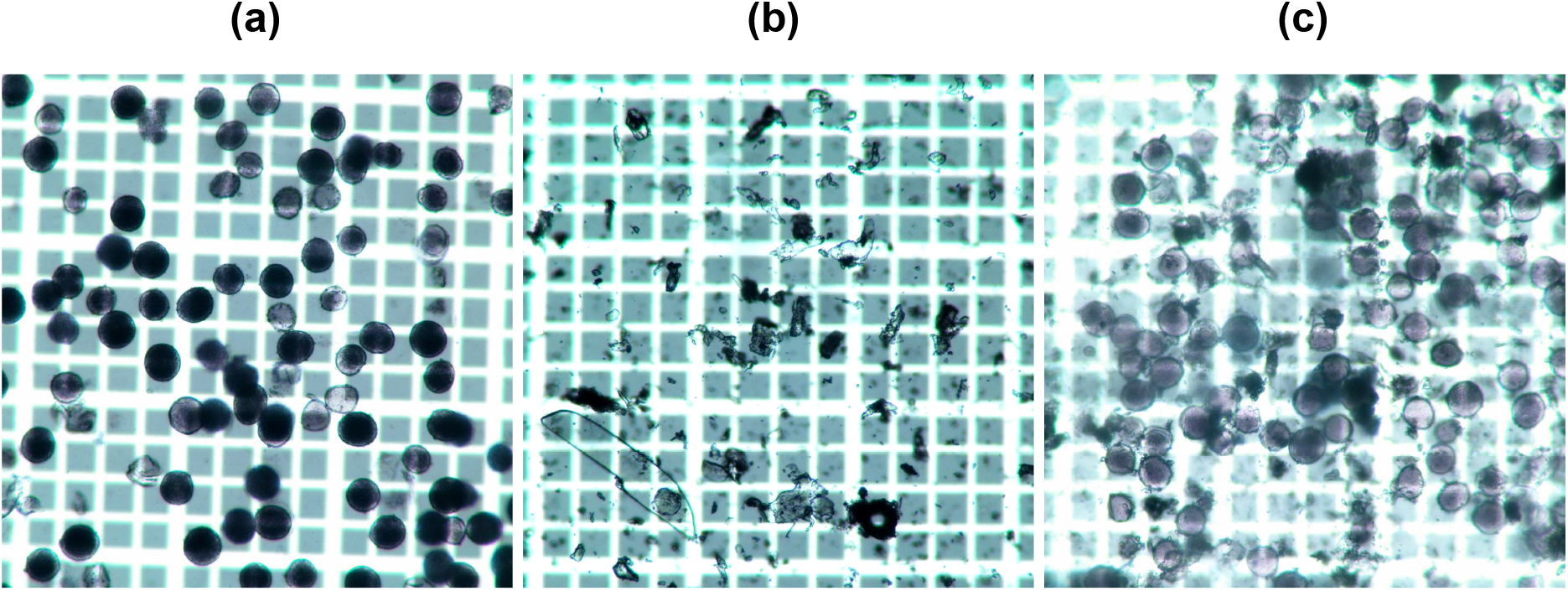
General aspect of **(a)** *E. lutea* pollen grains, **(b)** polipropilene fragments (<63 μm), and **(c)** a mix of pollen grains and plastic fragments. Images were obtained under light microscope (4x). Each small square unit is 25 μm.

**Figure S2.**
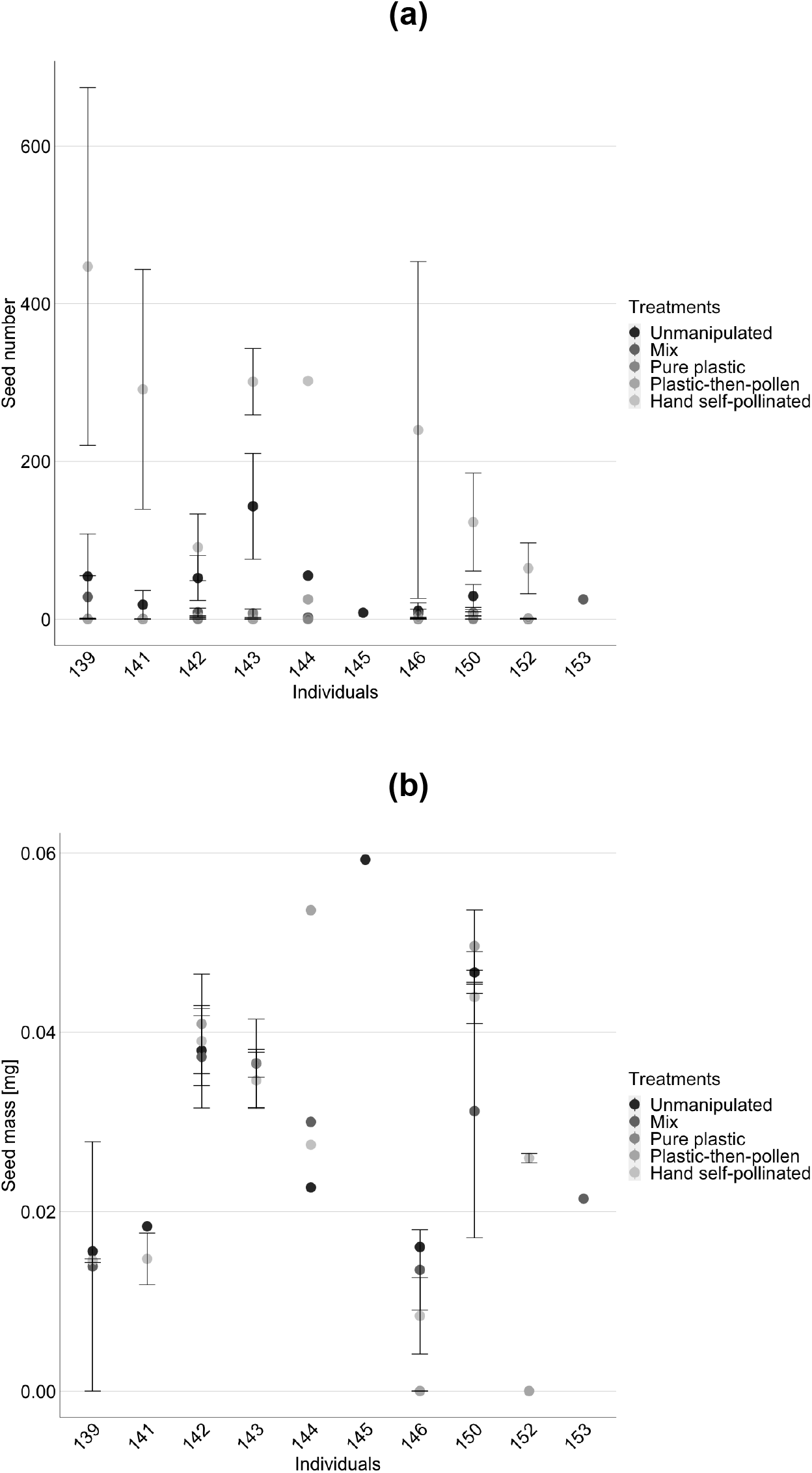
Effects of individual plants and pollination treatments on **(a)** seed number, and **(b)** mass per seed. Results of ANOVA-like table for random effects shown no statistical effects of individual plants on the seed number (*d*.*f*. = 1; *LRT* = 0.265; *P* = 0.606) but for seed mass (*d*.*f*. = 1; *LRT* = 26.31; *P*<0.001). *d*.*f*.: degrees of freedom for the likelihood ratio test; *LRT*: the likelihood ratio test.

**Figure S3.**
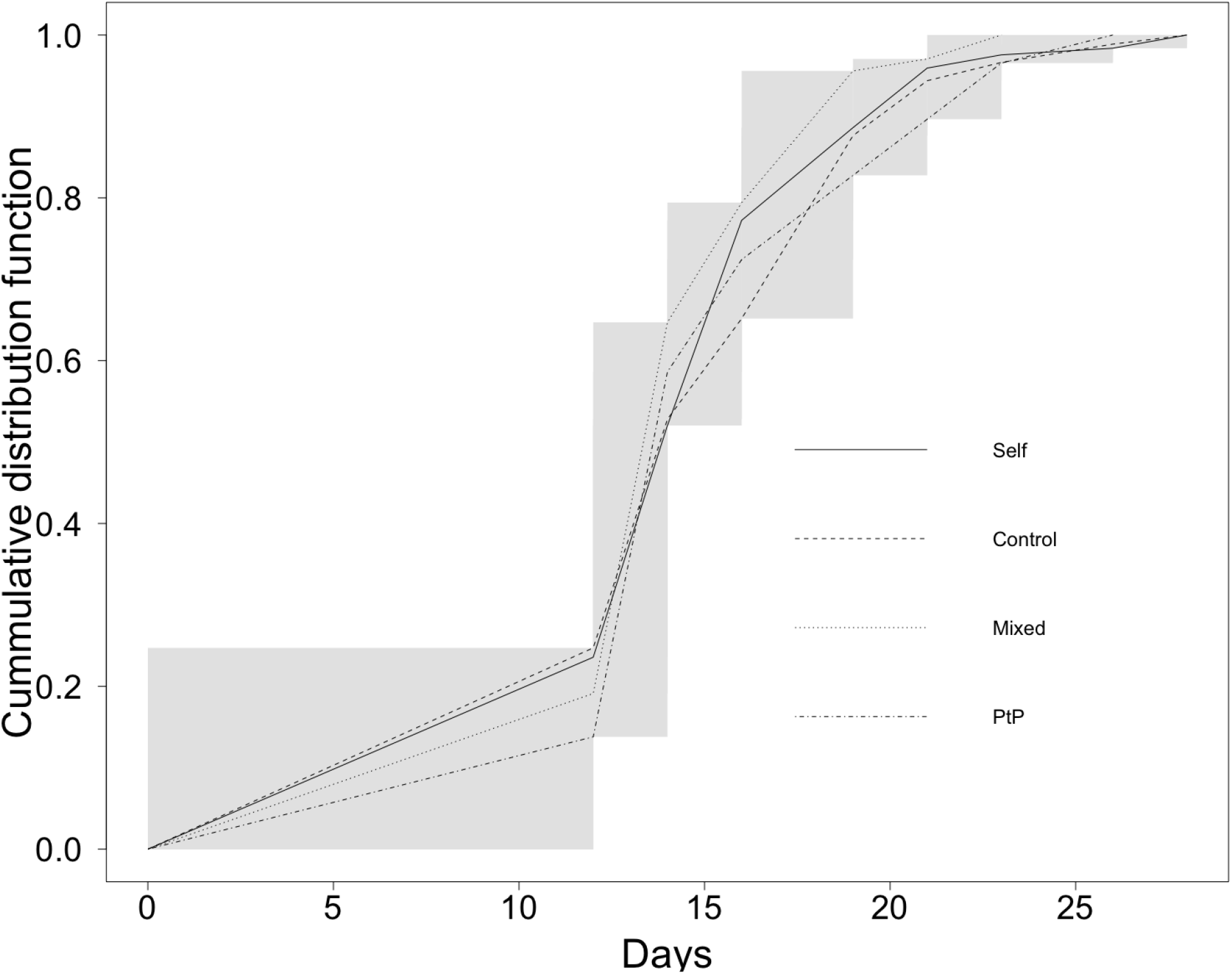
Cumulative distribution function of a nonparametric time-to-event curve for germination of seeds obtained under four pollination treatments: hand self-pollinated (Self), unmanipulated flowers (Control), polypropylene mixed with pollen (Mixed), and pollen deposited after to plastic deposit onto stigmas (PtP, pollen-then-plastic treatment). Gray areas represent the uncertainty due to censoring.

